# scGenAI: A generative AI platform with biological context embedding of multimodal features enhances single cell state classification

**DOI:** 10.1101/2025.05.07.652733

**Authors:** Ruijia Wang, Matthew Ung, Huanying Gary Ge

## Abstract

**Summary:** Single-cell sequencing has advanced the understanding of cellular heterogeneity, yet traditional cell type annotation tools struggle with increasingly complex datasets and multimodal integration. Recent large language models (LLMs) offer improved accuracy but rely on pre-trained models with limited gene vocabularies and lack of biological contextualization. These constraints make it challenging to effectively fine-tune pre-trained models, especially for novel datasets, non-human species, or disease-specific studies where unique gene expression patterns are critical. To address these constraints, we developed scGenAI, an LLM-based tool that supports straightforward and flexible *de novo* training, allowing researchers to incorporate genomic and biofunctional contexts to improve accuracy and interpretability on the prediction of cell states. Here, we demonstrate that scGenAI outperforms conventional models in tasks such as cell type prediction and acute myeloid leukemia (AML) malignant cell states identification, making it a powerful tool for single-cell analysis workflows.

**Availability and implementation:** scGenAI (DOI: <u>10.5281/zenodo.14927611</u>) is distributed as an open-source Python package, with source code and comprehensive documentation accessible on GitHub at https://github.com/VOR-Quantitative-Biology/scGenAI.

## 1. Introduction

Single-cell sequencing has expanded our understanding of cellular heterogeneity, enabling detailed cell type characterization and insights into disease mechanisms (Lähnemann *et al*., 2020). Traditional approaches for cell type annotation have been pivotal in analyzing single-cell RNA sequencing (scRNA-seq) data but are limited when handling complex datasets, particularly those integrating multimodal measurements (Butler *et al*., 2018) or containing disease-specific features such as chromosomal aberrations, which are hallmarks of many cancers and genetic disorders (Sansregret and Swanton, 2017). Recent tools such as scGPT (Cui *et al*., 2024) and scBERT (Yang *et al*., 2022) advancements in large language models (LLMs) have demonstrated the potential to capture intricate biological relationships and improve annotation accuracy. However, most of these tools rely on pre-trained architectures, are constrained by limited gene vocabularies, and lack comprehensive genomic and bio-functional contexts, which restrict their adaptability to novel datasets, non-human species, and disease-specific studies. To address these challenges, we developed scGenAI, a modular tool that supports *de novo* training and enables users to incorporate detailed biological knowledge for the specific prediction of cell types and cell states using single cell sequencing data. scGenAI offers a flexible, biologically informed framework for robust and accurate single-cell analysis, advancing computational biology research.

## 2. Methods and Features

scGenAI comprises five core modules for single-cell multimodal analysis including data preprocessing, input embedding, model training and evaluation, fine-tuning, and prediction (Figure 1A and Supplementary Methods). scGenAI’s preprocessing module prepares single-cell NGS data by filtering genes expressed in fewer than a specified number of cells, followed by normalization and transformation (Supplementary Method). Tokenization was then applied via the *GeneExpressionTokenizer()*, which encodes gene IDs and expression levels for context embedding (Figure 1A). scGenAI offers multiple embedding strategies to manage sequence length limitations and incorporates genomic and functional contexts controlled by the *scGenAI*.*context()*. These include *random, genomic, biofunction*, and *gene list* embeddings, which can be adopted for different research scenarios (Supplementary Methods). The random method shuffles gene tokens into overlapping windows of fixed length, genomic context arranges genes by chromosomal coordinates, and biofunction context groups genes by biological relevance. Specified gene lists enable selection to enhance key genes in multi-omics or multimodal applications (Supplementary Methods). Among all the methods, random context not only served as the baseline but also as a practical alternative when other context embeddings are unavailable. The random approach provides a straightforward fallback option that benefits from the flexibility of the model architecture while maintaining robust performance (Supplementary Figure 1). Other designed embedding methods leverage the concept of context embedding, which captures intrinsic relationships among genes based on genomic positioning, biological regulatory interactions, or user-defined gene sets. These embeddings can further enhance model performance in predicting cell types and states by incorporating biologically relevant information into the training process (Supplementary Methods).

**Figure 1.**
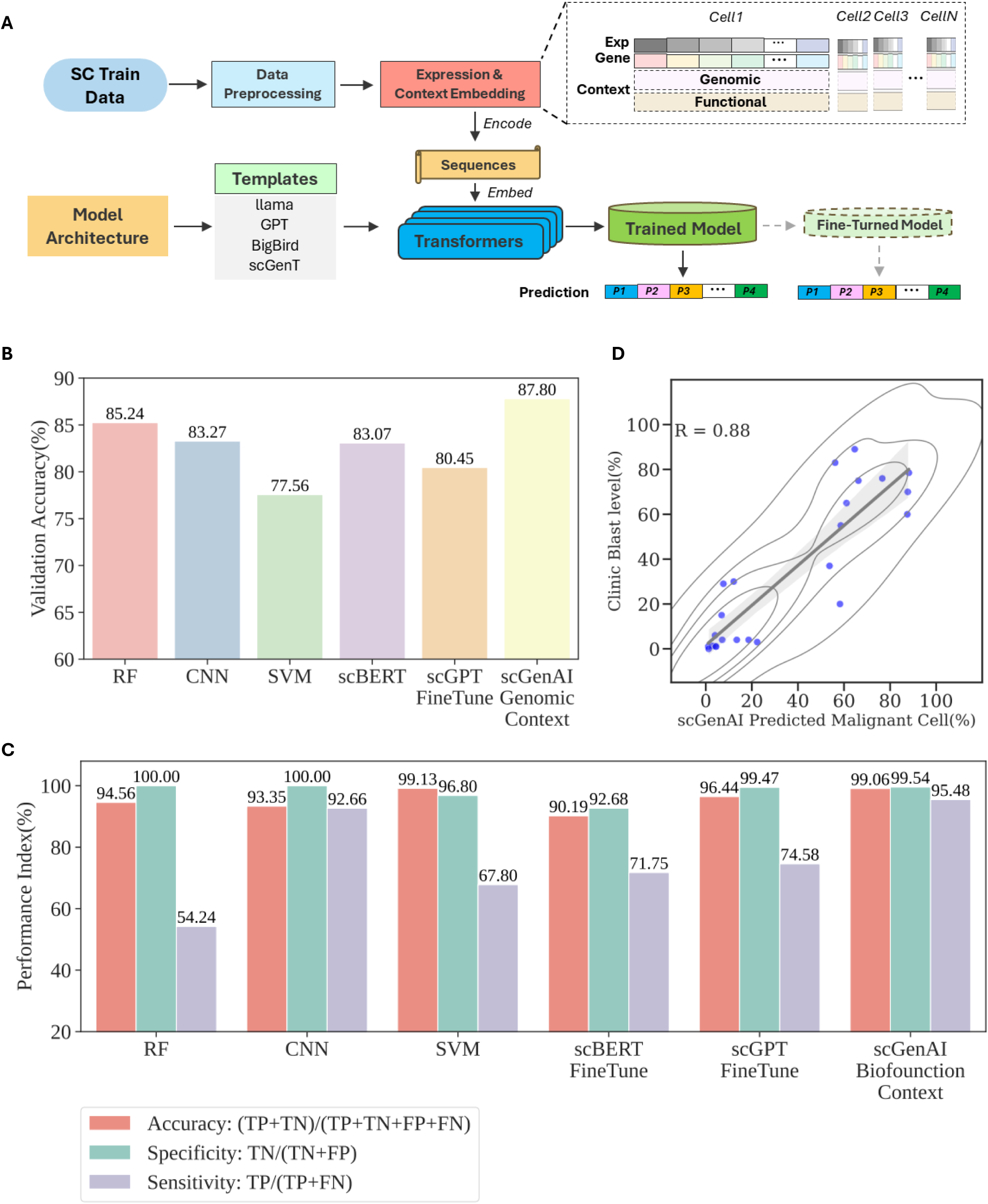
Design and Performance of scGenAI. **(A)** Schematic representation of the overall workflow and major modules in scGenAI. **(B)** Summary of cell type prediction accuracy for myeloid cells in myeloma tissue using scGenAI and other modeling methods. **(C)** Summary of model performance on AML malignant status using scGenAI and other modeling methods. **(D)** Scatter plot comparing the clinical blast level to the fraction of cells classified as malignant using scGenAI.

## 3. Application Examples and Results

### 3.1 Cell type model training and fine-tuning with multimodal single-cell data

To evaluate the performance of scGenAI in cell type annotation, we applied it to multiple datasets and compared its validation outcomes to other established models where applicable, including convolutional neural networks (CNN), random forests (RF), and support vector machines (SVM) as well as two LLM-based tools, scGPT (Cui *et al*., 2024) and scBERT (Yang *et al*., 2022) (Supplementary Methods). Two models were initially trained *de novo* using two datasets: a cancer related dataset of myeloma myeloid cells (GSE154763) (Cheng *et al*., 2021) and a non-human species dataset using mouse eye cells (GSE135167) (Lehmann *et al*., 2020) (Supplementary Methods). The LLaMA template was chosen for both datasets, based on its superior baseline performance (Supplementary Figure 1). To model myeloma myeloid cell types, which include seven cell population subsets of plasmacytoid dendritic cells, conventional dendritic cells, monocytes, and macrophages, we applied scGenAI context embedding to examine how genomic and biofunctional knowledge impact model accuracy. By embedding cytoband-based gene annotations for genomic context and a cancer-related gene set for the biofunction context (Supplementary Methods), scGenAI achieved accuracy levels of 83.27% to 87.80%, with genomic context yielding the highest accuracy (Supplementary Figure 2). The biofunction context method, informed by cancer-associated genes, achieved an accuracy of 87.66%, closely approximating that of the genomic context approach (Supplementary Figure 2). The higher accuracy of context compared to random methods and other modeling approaches demonstrated the benefits of integrating genomic and functional gene information (Figure 1B and Table 1). For the mouse eye cell dataset, scGenAI displayed the advantage of *de novo* training as it does not rely on a pre-trained model constrained by species or gene vocabulary limitations. The model achieved accuracy ranging from 98.55% to 99.27%, with the gene list method (using the top 1,000 variable genes) providing the highest accuracy. This was slightly higher than that of the models trained using the other methods (Table 1 and Supplementary Figure 2).

**Table 1.** Model performance (validation accuracy%) on scGenAI and other methods.

| Dataset/Method | RF | CNN | SVM | scBERT | scGPT | scGenAI |
| --- | --- | --- | --- | --- | --- | --- |
| Myeloid Cells | 85.24 | 83.27 | 77.56 | 83.07 | 80.45 | 87.80 |
| Mouse Eye Cells | 98.22 | 99.21 | 98.41 | N/A | N/A | 99.27 |
| Bone Marrow Fine-Tune | N/A | 76.78 | N/A | 79.80 | 78.93 | 79.90 |
| PBMC CITE-Seq | 86.93 | 85.46 | 85.42 | N/A | N/A | 87.50 |
| PMBC scATAC-Seq | 78.94 | 78.78 | 83.60 | N/A | N/A | 89.07 |
| AML Cells | 94.56 | 93.35 | 99.13 | 90.19 | 96.44 | 99.06 |

We also performed fine-tuned model training using two bone marrow datasets, GSE181989 (Tonglin *et al*., 2022) and GSE135194 (Wu *et al*., 2020), which present both common and unique cell types and expressed genes. scGenAI was first trained on GSE181989, achieving an accuracy of 79.90% using the random context method. The model was subsequently fine-tuned using GSE135194, which introduced two additional cell types and 3,368 newly expressed genes. After fine-tuning, the updated model successfully identified all cell types, with an accuracy improvement from less than 60% to 79.40%.

To evaluate the application of scGenAI to multimodal data, we used a peripheral blood mononuclear cell (PBMC) dataset containing both gene expression and surface antigen feature barcoding (CITE-seq) data (GSE164378) (Hao *et al*., 2021). We used scGenAI to train models on 17 cell types across different context methods with an emphasis on surface antigen expression. scGenAI outperformed other modeling methods and achieved a maximum accuracy of 87.50% by using a blood-related gene set with the biofunction context method (Table 1 and Supplementary Figure 2). Similar performance is also observed on another layer of multiomics data using a PBMC scATAC-Seq dataset (Stuart *et al*., 2019), in which scGenAI achieved a maximum accuracy of 89.07% (Table 1 and Supplementary Figure 2).

### 3.2 AML Malignant Cells Prediction

Beyond standard cell type annotation, we evaluated scGenAI’s capability to classify malignant cells within a mixed population of leukemic and healthy cells in a multimodal dataset of acute myeloid leukemia (AML) (GSE116256) (van Galen *et al*., 2019) (Supplementary Method). While both healthy cell type annotation and malignant cell state prediction fall under the broader category of cell annotation, identifying malignant cells is inherently more complex. Malignant cell status is typically determined through mutation calling from sequencing data and is often accompanied by distinct mRNA expression patterns (van Galen *et al*., 2019).

Model performance was evaluated by measuring the accuracy of malignant status prediction alone and in combination with cell type classification. While all models exhibited similar specificity levels, scGenAI and SVM exhibited higher accuracy (∼99%) in predicting malignant states, with scGenAI achieving the highest sensitivity (95.48%) among both traditional regression methods and pre-trained models including scGPT and scBERT (Figure 1C and Supplementary Methods). Given the limited representation of malignant cells (12%) in the training and validation datasets, scGenAI’s superior sensitivity and specificity highlight its robustness in accurately predicting cell states, both for broad cell types determined by marker gene expression and more opaque states that arise from gene mutations. Notably, the use of cancer-related gene sets in scGenAI’s context displayed the highest combined accuracy (81.80%) for both cell type and malignant status, outperforming all other methods tested (Supplementary Figure 3). Furthermore, the scGenAI-trained model was applied to the entire AML dataset to estimate the percentage of AML blasts in each sample. These predictions were compared with percentages obtained from flow cytometric sorting. The blast percentages predicted by the scGenAI-trained model showed a high correlation with the experimental results (R = 0.88) (Figure 1D). This strong correlation confirms scGenAI’s effectiveness in accurately identifying AML blasts within complex, mixed-cell populations.

## 4. Conclusions

We introduced scGenAI, a large-scale model-based tool to address the key limitations of existing tools by enabling training from *de novo* and incorporating genomic and biofunctional contexts. Our results demonstrate that scGenAI not only delivers robust performance in predicting cell types and malignant cell states across both single and multimodal datasets, but also allows researchers to train models *de novo* using their own high-quality internal datasets, establishing it as a valuable resource for advancing single-cell states prediction across diverse applications.

## Supporting information

Supplementary Infomation

## Author’s Contributions

R.W., M.U, and G.G. conceived of and designed the program and wrote the paper. R.W. implemented the method and analyzed the data.

## Acknowledgments

We thank the members of Vor Bio’s Research Department for their helpful discussions.

