## Supplementary Figures for "scGenAI: A generative AI platform with biological context embedding of multimodal features enhances single cell state classification"

**Supplementary Figure 1**

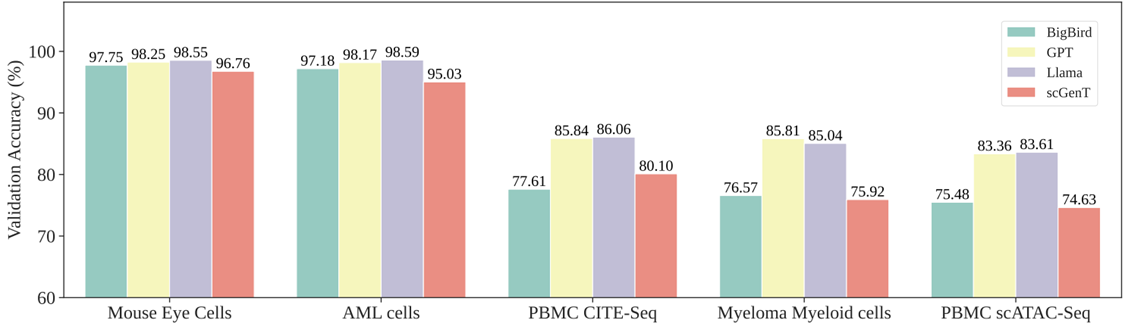

**Supplementary Figure 1** Accuracy summary of the model template in scGenAI using random context method

**Supplementary Figure 2**

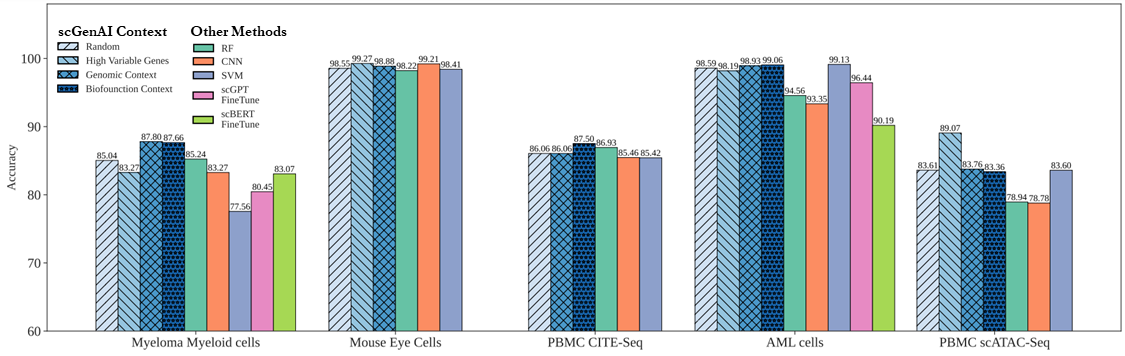

**Supplementary Figure 2** Summary of scGenAI training across datasets.

Model accuracy achieved using different context embedding methods in scGenAI as well as other modeling methods across each training dataset. scGPT and scBERT model fine-tuning are not evaluated in the three datasets including mouse eye cels, CITE-Seq and scATAC-Seq due to their limitation of pre-trained models which relies on human gene expressions.

**Supplementary Figure 3**

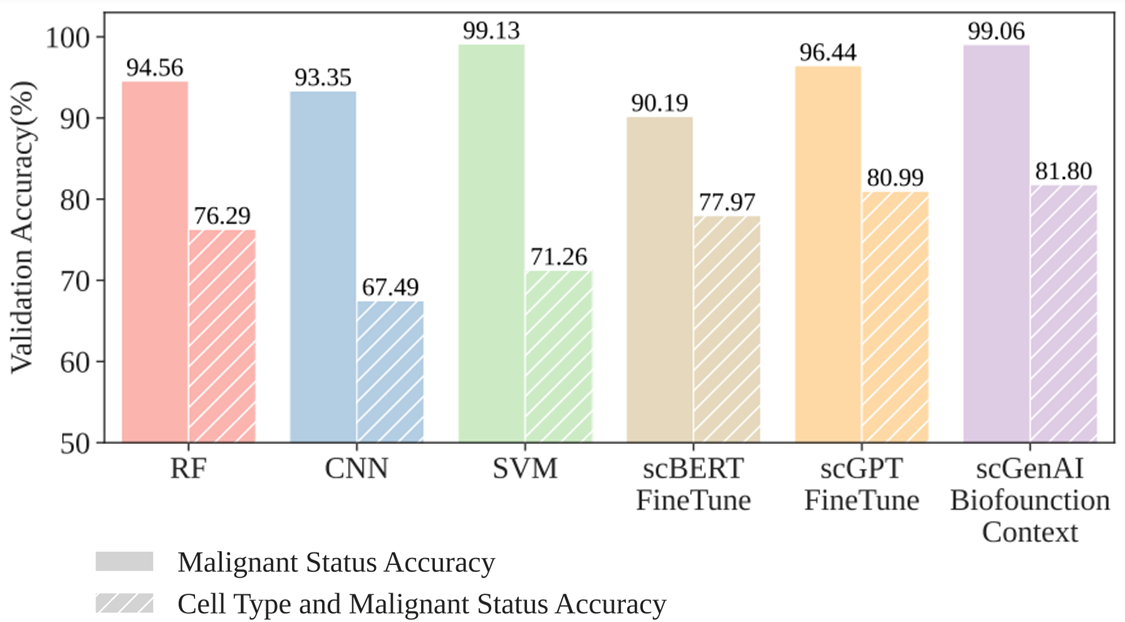

**Supplementary Figure 3** Summary of cell type and AML malignant status accuracy using scGenAI and other modeling methods.

**Supplementary Figure 4**

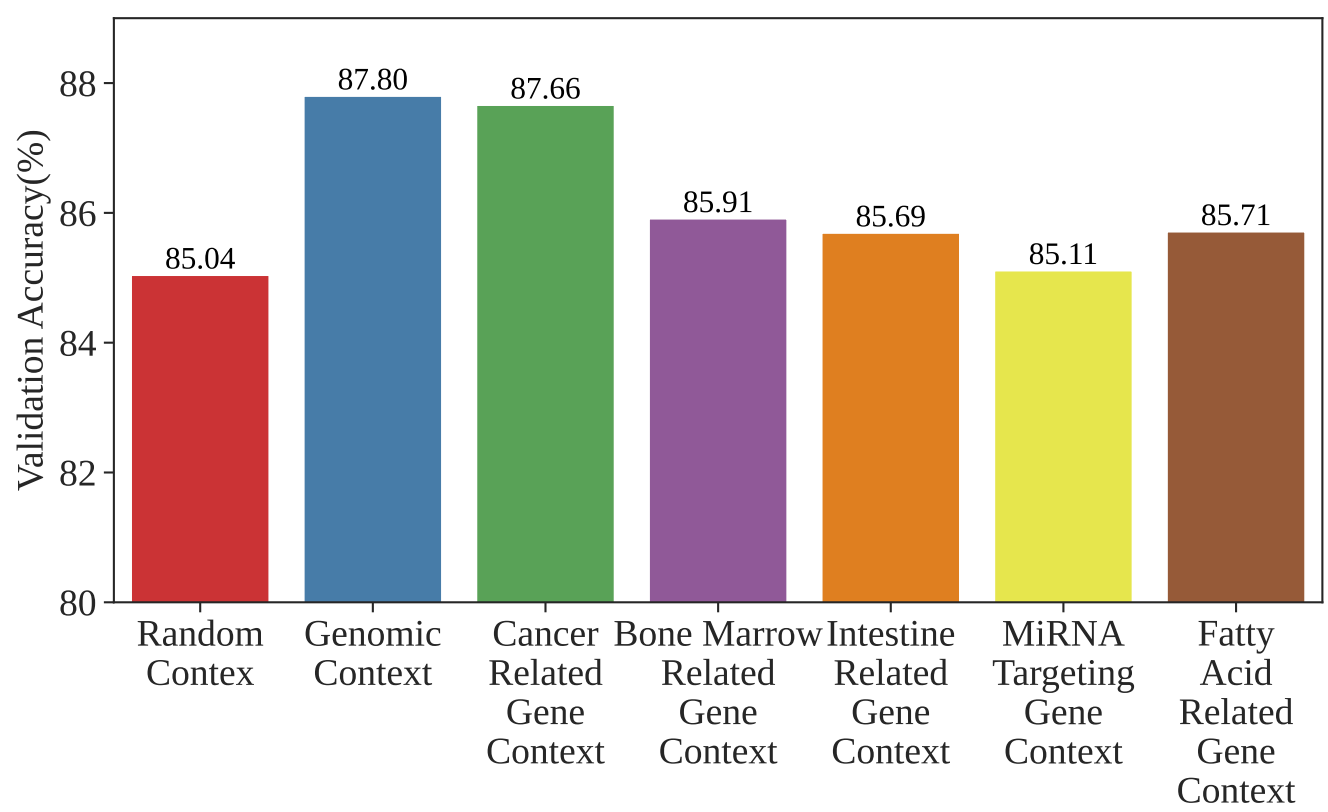

**Supplementary Figure 4** Summary of cell type prediction accuracy using different context methods on the Myeloma Myeloid cells dataset.
