## Supplementary Infomation for "scGenAI: A generative AI platform with biological context embedding of multimodal features enhances single cell state classification"

**Supplementary Methods Information**

**Preprocessing Module in scGenAI**

Prior to model training, single-cell NGS data were preprocessed using the “*Preprocessor*” class. Genes expressed in fewer than a defined number of cells (default of 50) were removed to eliminate low-quality and uninformative features. The remaining genes were normalized to a total count of 10,000, followed by ln(X + 1) transformation. The transformed expression values were discretized into bins (default 10) based on their distribution within each cell.

**Knowledge context embedding in scGenAI**

scGenAI uses several methods to fragment sequences, address sequence length limitations, and incorporate gene functions and genomic contexts. These strategies help to capture key biological features, such as cell type, cell genotype, and cell state.

*Random context approach*

The random approach shuffles genes and divides each tokenized sequence into an overlapping context of fixed length. A specified stride, defaulting to half the context length, dictates the step size between the contexts. Padding tokens were added to ensure consistent context length. Among all the methods, the random context embedding serves as the baseline procedure within scGenAI. In scenarios where users need to train a single-cell model from scratch but lack an appropriate genomic or biofunctional context, random embedding provides a straightforward default option.

*Genomic context embedding approach*

The Genomic embedding function generates the context of gene expression data based on genomic coordinated groups (e.g., cytobands). Genes were grouped by cytoband, combined into sequences, and repeated to a specified depth (default 2). These sequences are then divided into contexts based on cytoband boundaries closest to the desired length.

*Biofunctional or genic functional context embedding approach*

scGenAI also embeds gene expression data in the context of gene sets and pathways. Gene sets were filtered to retain only expressed genes, and a random sampling approach generated contexts from the same gene set or pathway. Sampling was iterated until each gene set was sampled to a specified depth (default 2). Genes not included in the set were context using the random method.

*Gene set and manual selection approach*

scGenAI offers options to either subset data using a gene list or to enhance specific genes. In the subset approach, the dataset is reduced to the selected genes and contextualized using the random sliding method. The emphasis approach duplicates key genes (or surface antigens) from a predefined list within a sequence using an emphasis factor. These duplicates were inserted randomly to avoid positional bias, which is particularly useful for single-cell multi-omics data such as CITE-seq.

*Usage of different context approaches*

All the embedding methods introduced in scGenAI are to capture underlying relationships among genes, employing genomic, biofunctional, or user-defined gene sets that may be relevant for predicting cell types and states. For instance, genomic context embeddings can help uncover phenotypic differences linked to chromosomal instability (CIN), a feature of critical importance in understanding cancer progression (Burrell et al., 2013). Biofunctional context embeddings, on the other hand, leverage known gene pathways or networks (Tirosh et al., 2016), while the gene list context facilitates analyses focused on a specific subset of genes (e.g., top variable genes). In scenarios where a suitable biological or genomic context is not available, random embeddings provide a straightforward fallback option, one that nevertheless benefits from the GPT/LLama-like model architecture and demonstrates robust performance (Supplementary Figure 1). Each of these embedding approaches has distinct advantages and constraints. Genomic and biofunctional embeddings can offer greater biological interpretability but may require more domain-specific knowledge, whereas random embeddings demand minimal prior information. Comparing to text-based strategies like scBERT (Yang *et al.*, 2022), which impose an artificial ordering on gene expression data, scGenAI emphasizes biologically informed relationships that might be less evident under text-only encodings. When users have information about genomic positional or functional gene sets relevant to their single-cell dataset, we recommend employing either the genomic or biofunction embedding methods. For instance, utilizing cytoband-based gene sets or cancer-related gene sets in tumor datasets. If users are uncertain about which gene set to select, we suggest starting with simpler embedding strategies, such as random context or a gene list embedding that focuses on top variable genes. As shown in Supplementary Figure 4, using an irrelevant gene set does not improve performance compared to the baseline random context. All the genomic context file and biological context gene set files used in this study is downloaded from the Molecular Signatures Database (MSigDB, v2023.2) (Liberzon *et al.*, 2011).

Once the context is generated, tokenized sequences, labels, and positional encodings are loaded into *DataLoader* instances to batch the data for efficient GPU utilization.

**Dataset Processing for Training and Validation**

Several datasets were used to assess the *de novo* training performance of scGenAI, including myeloid cells (GSE154763), mouse eye cells (GSE135167), and AML cells (GSE116256). Each dataset was randomly split into training and validation sets at a 4:1 ratio, ensuring a proportional representation of cell types across both sets. For AML malignant cell state identification, which is another layer of cell annotation, only cells confirmed to harbor AML-related mutations thought nanopore sequencing were retained as training targets.

To evaluate the fine-tuning capabilities of scGenAI, two bone marrow datasets, GSE181989 and GSE135194, were used. The training and validation cells were split similarly, with GSE181989 data used for the initial round of training, followed by GSE135194 for the fine-tuning phase. The validation datasets from both GSE181989 and GSE135194 were combined to assess the fine-tuned model performance at each checkpoint.

For multi-omics data analysis, the PBMC CITE-Seq dataset (GSE164378) containing both RNA and surface antibody abundance profiles was employed. The analysis focused on sequencing lane 1, which included 7,619 cells across 17 annotated cell types, including B, myeloid, NK, and T cells. The dataset was processed and split in the same manner as the other scRNA-Seq datasets, allowing consistent training and validation procedures.

The AML cells dataset contained single-cell mutation calling information, which we used as the ground-truth label for malignant cells. The model was trained on 7,444 out of 38,410 cells from 40 bone marrow aspirates, including samples from 16 AML patients and five healthy donors (van Galen et al., 2019). The model was trained using labels that combined both cell type and malignant status

All training and validation datasets used in this study have been deposited in Zenodo (<https://zenodo.org/>) and are accessible through Zenodo ID **14036273, 14036352, 14036400, 14036452, 14036508,** and [**14867904**](https://doi.org/10.5281/zenodo.14867904)**.**

**Model Templates in scGenAI**

The scGenAI framework integrates multiple model templates to support effective sequence classification in single-cell multi-omics data. These templates, which include LLaMA, scGenT, GPT, and BigBird-based architectures, provide a customized classification layer that is capable of predicting cell types, genotypes, and disease states from complex input data.

The *CustomLLaMAForSequenceClassification* model templates leverage the LLaMA architecture, which was initially developed to perform sequence classification for causal language modeling. The model integrates *LlamaConfig*, enabling the customization of key parameters such as hidden size and maximum sequence length, which permits flexibility in handling variable input dimensions. The classification layer is designed to be dynamically adjustable by matching the hidden state dimensions; when the hidden dimensions change, as during fine-tuning, the classifier layer adapts accordingly. The embedding component supports token resizing to accommodate changes in gene or antibody token counts during training or fine-tuning, thereby facilitating a broader scope of data input.

The *scGenT* model template, *CustomscGenTForSequenceClassification*, is tailored for tasks involving single-cell data. Its byte-pair encoding, multi-head attention, and layer normalization provide robust handling of the gene expression sequences. The attention mechanism uses a custom scGenT attention mechanism with a triangular mask that preserves gene sequence ordering and effectively captures the dependencies among genes. Additionally, prelayer normalization was applied to improve the model stability and convergence, ensuring smoother gradients across the transformer layers.

The GPT-based template within scGenAI, *CustomGPT2ForSequenceClassification*, adapts the GPT-2 model for sequence classification tasks, allowing the flexible resizing of vocabulary and hidden layers to meet the demands of specific single-cell contexts. The model incorporated a linear classification layer to process the final hidden state of each token sequence, yielding logits with adjustable dimensions to match the training requirements. The *GPT2Config* configuration manages key model parameters, including the embedding size, attention heads, and number of layers, allowing for optimized configurations across diverse single-cell datasets.

The *CustomBigBirdForSequenceClassification* template, which is based on BigBird, is optimized for longer sequences, making it particularly suited to high-dimensional single-cell data. BigBird’s sparse attention mechanism enables the efficient processing of extended sequences, which is especially useful in high-dimensional genomics datasets. In addition, adjustable dropout parameters, including *hidden_dropout_prob* and *attention_probs_dropout_prob*, control overfitting and enhance model robustness, enabling stable performance across large input sizes.

Each model template utilizes a specific initializer class to configure and load its tokenizer, thereby facilitating targeted processing of single-cell data. *BigBirdModelInitializer* and *GPTModelInitializer* load the respective tokenizers for BigBird and GPT, supporting unique tokenization schemes for gene and expression data. These initializer classes also allow adjustments to parameters such as vocabulary size, hidden dimensions, layer numbers, and sequence length, ensuring compatibility with a wide range of single-cell datasets.

**Customized Model Architecture and Training**

scGenAI supports customizable transformer-based model architectures via the *scGenAI.Models()*, with templates such as LLaMA, GPT, BigBird, and scGenT. Each architecture combines a transformer with a classification head, enabling the cell type, cell state, or genotype prediction. The transformer processes the input sequences to generate hidden states that capture complex expression patterns. A linear classification layer applied to the final hidden state produces logits for prediction. Model optimization is guided by cross-entropy loss, with the performance assessed using *model_train_and_eval()*. Checkpoints are saved based on the best validation accuracy and final epoch performance.

**Prediction and Fine-tuning Module in scGenAI**

In the prediction module, the datasets were processed using the same tokenization and context embedding as in the training phase. The *Prediction()* function assigns the most probable class per context and then aggregates predictions across multiple contexts per cell to provide the final prediction.

When the fine-tuning module is selected, the pre-trained models can be updated using new datasets. After loading the model files and fine-tuning the dataset, scGenAI regenerates contexts based on the pre-trained model configuration. Gene vocabulary and training features (e.g., cell types) were compared between the pre-trained model and the new dataset, and updated as necessary. The model was retrained, with adjusted weights, and saved as a refined model for future use.

**Benchmarking with Public Tools**

To evaluate scGenAI’s performance, we benchmarked it against several established machine learning models—convolutional neural networks (CNN), random forests (RF), and support vector machines (SVM)—implemented using the Python scikit-learn and PyTorch libraries. All models were trained on the same preprocessed datasets to ensure a standardized and fair comparison. Pearson’s correlation between predicted malignant cell percentage using AML dataset and experimental blast percentage of each sample is calculated through SciPy. Additionally, scGPT (Cui *et al.*, 2024) and scBERT (Yang *et al.*, 2022), large language model (LLM) tools, were included for benchmarking. The analysis of fine-tune using bone marrow was not performed for RF and SVM. For *scGPT* and *scBERT*, model fine-tuning was conducted using their pre-trained, human-specific models on human datasets only, as neither tool currently supports training from scratch using non-human genes. Additionally, due to the absence of specific documentation for handling multi-omic datasets, CITE-seq and scATAC-Seq could not be tested using *scGPT*’s and *scBERT’s* fine-tuning protocol. Comparative performance was assessed by measuring accuracy on the validation sets to evaluate *scGenAI*’s robustness and accuracy relative to both traditional methods and LLM-based approaches.

**Computational Resource**

All analyses in this study were conducted on an AWS *g4dn.12xlarge* instance, which provides 48 CPUs, 192 GB of memory, and 4 GPUs, each with 16 GB of memory (totaling 64 GB GPU memory).
